# Methamphetamine modulates functional connectivity signatures of sustained attention and arousal

**DOI:** 10.1101/2025.05.20.655181

**Authors:** Yizhou Lyu, Anna Corriveau, Hanna Molla, Harriet de Wit, Monica D. Rosenberg

## Abstract

Building on evidence that psychostimulants modulate whole-brain functional connectivity signatures of sustained attention, we examined how a single dose of methamphetamine (MA, 20 mg) changes network-level functional organization and sustained attention in healthy adults. Using a within-subject, placebo-controlled design, we tested whether MA selectively affects behavioral and fMRI connectivity signatures of sustained attention and arousal. Under MA, participants showed improved sustained attention task performance as well as functional connectivity signatures of higher sustained attention and arousal. These network changes emerged consistently across resting-state and task-based fMRI, indicating that MA influences attention- and arousal-related networks regardless of cognitive context. Furthermore, a support vector classifier distinguished functional connectivity patterns observed during the MA and placebo conditions, identifying connections overlapping with networks related to arousal. Together, these findings align with prior work on other psychostimulants like methylphenidate, showing that MA modulates sustained attention and related large-scale brain networks. By revealing how MA modulates attention-relevant brain connectivity patterns, our results highlight the utility of psychostimulants as causal tools for probing the robustness, generalizability, and interpretability of brain-based biomarkers of behavior.

## Introduction

Attention is a fundamental cognitive process that can be modulated by psychostimulant medications. Sustained attention, the ability to maintain focus on a task over time, is both a stable trait and a dynamic state that recruits networks across the brain (Rosenberg et al., 2016a). Sustained attentional performance can be enhanced or impaired by factors such as cognitive conditions and pharmacological interventions (Swanson et al., 2011). Indeed, pharmacological stimulants are commonly prescribed for the treatment of disordered sustained attention, such as attention-deficit/hyperactivity disorder (ADHD). What are the effects of these medications on functional brain networks known to be involved in sustained attention? In the current study, we test the effects of a dose (20mg) of the psychostimulant methamphetamine (MA) on predefined attentional networks during fMRI tasks and rest and on behavioral performance during a sustained attention task.

Methamphetamine, like methylphenidate and d-amphetamine, is approved as a treatment for attention-related disorders. In addition to their use as treatments for attention-related disorders such as ADHD (Faraone, 2018), amphetamines are known to enhance attention and cognitive control. Amphetamines such as methamphetamine and d-amphetamine increase the synaptic levels of dopamine and norepinephrine in the brain by releasing presynaptic stores of the neurotransmitters, blocking reuptake (Avelar et al., 2013; Covey et al., 2013), and inhibiting their breakdown by monoamine oxidase (Riddle et al., 2007; Miller et al., 1980). These actions strengthen signaling pathways in critical areas for attention and executive processing, such as the prefrontal cortex and striatum (Faraone et al., 2015). Neuroimaging studies have found that amphetamine produces global changes in the brain, including altering whole brain cerebral blood flow to areas of the brain with dopamine innervation, such as in the striatum, anterior cingulate cortex, prefrontal and parietal cortex, inferior orbital cortex, thalamus, cerebellum, and amygdala (Devous et al., 2000; Rose et al., 2006; Vollenweider et al., 1998). Functional MRI studies in healthy adults have also shown that amphetamine induces changes in BOLD signal variability in prefrontal and subcortical regions, along with improvements in working memory task performance, which is related to attention (Garrett et al., 2015; Crane et al., 2023). Repeated use of MA leads to disrupted functional brain organization and abnormal network interactions (Ahmadlou et al., 2013), and chronic MA users have abnormal connectivity between cognitive control networks and the default mode network, leading to executive functioning deficits (Ipser et al., 2018).

Methylphenidate, also used in the treatment of ADHD (Contini et al., 2013), elevates dopamine and norepinephrine levels mainly by blocking their re-uptake transporters, thereby prolonging the neurotransmitters’ signalling time (Volkow et al., 2001; Berridge et al., 2006; Spencer et al., 2015). In contrast, amphetamine-class drugs increase neurotransmitter levels by enhancing release from the presynaptic neuron (Faraone, 2018). Despite these mechanistic differences, amphetamine and methylphenidate have similar behavioral effects. They improve attention and psychomotor functioning (Silber et al., 2006) and cognition and executive function (Moeller et al., 2014), as well as changes in risky decision making (Oswald et al., 2015) and reward processing (Stoy et al., 2011; Wei et al., 2021).

Functional connectivity (FC), defined as the temporal correlation between neuronal activation in spatially distinct brain regions, has become an important measurement in neuroimaging studies in the past decade (Van Den Heuvel & Hulshoff Pol, 2010). Whole-brain functional connectivity analyses using fMRI have identified biomarkers linked to distinct brain states, including sustained attention (Rosenberg et al., 2016a) and arousal (Ke et al., 2025), as well as individual phenotypes such as fluid intelligence (Finn et al., 2015). These connectivity-based measures enhance our understanding of how factors like drug administration can induce changes within specific brain networks. For example, previous work suggested that methylphenidate affects the sustained attention connectome-based predictive model (saCPM)—a functional connectivity neuromarker of sustained attention (Rosenberg et al., 2016a)—in predictable ways. Individuals given methylphenidate exhibited higher strength in the saCPM’s high-attention network and lower strength in the saCPM’s low-attention network compared to unmedicated controls (Rosenberg et al., 2016b). In contrast, individuals showed the opposite pattern—functional connectivity signatures of *worse* sustained attention in the saCPM’s constituent networks—when given the anesthetic agents propofol (Rosenberg et al., 2020; Chamberlain & Rosenberg, 2022) and sevoflurane (Rosenberg et al., 2020). Given common effects of amphetamine and methylphenidate on dopamine and norepinephrine modulation, we hypothesize that, similarly to methylphenidate, MA will produce similar selective changes in attention-related brain networks. Specifically, we employ the saCPM and a recently developed arousal CPM to assess MA’s influence on attention-related processes. We anticipate effects of MA on both arousal, defined as a state of physiological alertness and activation, and sustained attention, the ability to maintain focus on a task at hand (Esterman & Rothlein, 2019).

In the present study, we examine whether pharmacologically induced changes in cognitive and attentional state are reflected in functional connectivity patterns. We compare the strength of networks predicting sustained attention and arousal in healthy adults given a single dose of MA with network strength in a placebo condition during both resting-state and task-based fMRI. Participants under the MA condition demonstrated increased high attention and arousal network strength and decreased low attention and arousal network strength relative to the placebo condition, as observed during both resting-state and task-based fMRI. Additionally, we test whether these changes in functional connectivity correspond to changes in sustained attention task performance and whether the effects differ between individuals with varying baseline levels of attention. Our findings aim to clarify the neural mechanisms underlying MA’s effects on attention and contribute to our understanding of how psychostimulants modulate whole-brain functional connectivity networks. More broadly, our work emphasizes the value of psychostimulants as causal manipulation tools that offer a new kind of opportunity to test the robustness and generalizability of behavioral neuromarkers.

## Materials and Methods

### Design

The study used a within-subject design in which healthy adults participated in two MRI and two behavioral laboratory sessions during which they received MA (20 mg) or placebo. During scan sessions, functional MRI data were collected at the time of peak effect. During behavioral sessions, participants performed a sustained attention task outside the scanner. Participants completed self-report questionnaires during the sessions. The study was approved by the University of Chicago Institutional Review Board.

### Participants

We used a dataset collected and described in previous work (Molla et al., 2023; Molla et al., 2025). Briefly, healthy male and female volunteers (n = 104) between the ages of 18 and 35 were recruited from nearby college campuses and surrounding communities through flyers and advertisements on social media. Participants underwent in-person screening including a physical exam, electrocardiogram, psychiatric screening interview, medical, and drug use history assessments. Inclusion criteria included being right-handed; normal electrocardiogram; fluent in English; body mass index between 19 and 26; and at least a high school education. Exclusion criteria were history of psychosis; evere post-traumatic stress disorder; depression; current suicidal ideation; prescription medication use (other than birth control); contraindications for MRI scan; pregnancy (verified by pregnancy tests on sessions days); history of cardiovascular disease; or consuming more than 4 alcohol or caffeinated beverages a day. Females who were not on hormonal birth control attended sessions during the follicular phase of their menstrual cycle.

### Study procedure

Participants first attended a screening and orientation session, where they provided informed consent and were familiarized with the study procedures, and completed the pre-study gradual-onset continuous performance task (gradCPT). During this visit, they were introduced to the tasks they would complete throughout the study. After this visit, eligible participants took part in two behavioral sessions in which they received oral methamphetamine (MA; 20 mg) and placebo (PL) in a randomized, double-blind manner. During the behavioral sessions, participants completed mood and drug effect questionnaires, along with various cognitive tasks including the gradCPT. Of these data, only gradCPT performance is analyzed here.

Following the behavioral sessions, participants completed two fMRI scan sessions, also involving randomized administration of MA and placebo. To control session time and avoid participant fatigue, the behavioral measurements and fMRI scans were conducted on two separate drug administration days, thereby keeping each session at a reasonable length. We assume that the same participant would experience similar effects from the same drug administered on different days. This resulted in 5 study visits total (one screening/orientation visit, two behavioral visits, and then two MRI visits).

In the fMRI sessions, participants completed runs in the following order: a Monetary Incentive Delay (MID) task run (Knutson et al., 2000), two Doors task runs (Carlson et al., 2011), a 7-minute resting-state run (except for the first two participants, who had an 8-minute rest run), and an arterial spin labeling (ASL) run. To ensure effects were due to the experimental drug manipulation, participants were required to refrain from recreational drug use for at least 48 hours and from consuming alcohol for 24 hours before each session. Abstinence was verified through self-reported data, breath alcohol analysis, and urine drug tests.

During MRI sessions, participants were administered either MA (20 mg) or a placebo in randomized order, with each session occurring from 9:00 AM to 1:00 PM and spaced at least 72 hours apart (Molla et al., 2023). Participants were instructed to fast for eight hours prior to each session, after which they received a light snack. They were informed that they might receive a stimulant, sedative, or placebo. During the sessions, participants completed baseline measures for subjective and cardiovascular ratings before receiving either the MA or placebo at 9:30 AM. MA (Desoxyn tablets with dextrose filler) or placebo (dextrose) was administered in opaque capsules under double-blind conditions. Following drug administration, participants completed several subjective assessments, including drug effect questionnaires, at multiple time points.

Approximately 70 minutes after taking the capsule, participants were escorted to undergo the MRI scan. Throughout the session, cardiovascular measures were monitored, and sessions ended when participants’ blood pressure and heart rate was within 20% of baseline. At the end of all sessions, participants were debriefed and compensated for their time.

### Behavioral gradual-onset continuous performance task

The gradual-onset continuous performance task (gradCPT) was employed to assess sustained attention and inhibitory control (Esterman et al., 2013) outside the MRI scanner. The stimuli consisted of grayscale images of cityscapes and mountain scenes. During each 800 ms trial, one image transitioned gradually to the next through pixel-by-pixel linear interpolation. Participants were instructed to press a button in response to city scenes (90% of trials) and to withhold their response to mountain scenes (10% of trials). Participants completed 750 trials (10 minutes) per session. Task performance was evaluated using sensitivity (*d’*), the difference between standardized hit and false alarm rates.

### fMRI Tasks

#### Doors task

During fMRI, participants completed the “Doors” task, a reward-guessing game designed to assess an index of reactivity to monetary rewards and losses. The task was adapted from prior studies (Crane et al., 2023). In each trial, participants used a button box to choose between two doors, one hiding a reward of $0.50 and the other a loss of $0.25. Participants were told their performance could earn them up to $15. However, outcomes were predetermined, with 30 wins and 30 losses presented in a pseudorandom order across two runs. The task lasted 15 minutes total, split into two runs of approximately 7.5 minutes each, collected back to back with a short break in between. To ensure task engagement, participants were excluded if they responded to fewer than 75% of trials over the two runs, consistent with the procedure used in previous studies (Crane et al., 2023).

#### Monetary Incentive Delay (MID) task

The Monetary Incentive Delay (MID) task (Knutson et al., 2000) was used to elicit neural responses during both anticipation and receipt of monetary gains and losses. At the start of each trial, participants viewed one of six cues (circle for gains, square for losses) for 2,000 ms, signaling the potential of win or loss. Specifically, the task included two valences (gain vs. loss) and three magnitudes ($5, $1, $0) conditions, resulting in six total conditions (±$5.00, ±$1.00, ±$0.00). Following the cue, a fixation cross appeared for 2,000–2,500 ms (anticipation phase) and then participants were shown a triangle target (150–500 ms) and instructed to press a key before it disappeared. Success resulted in either a monetary gain or avoiding a loss, depending on the trial type. Feedback about the outcome was provided for 2,000 ms. Trials were separated by an intertrial interval of 2,000–6,000 ms, with a total of 90 trials (15 per condition) presented in randomized order. The full task lasted approximately 18.5 minutes. The target duration was adaptively adjusted to maintain a 66% hit rate, and participants with hit rates below 44% were excluded (replicating Molla et al, 2023).

### fMRI data acquisition

Functional MRI scans were conducted at the University of Chicago Magnetic Resonance Imaging Research Center on a 3 T Philips Achieva scanner with a 32-channel head coil. Data were collected using a gradient-echo echo-planar imaging (EPI) sequence with the following parameters: TR = 2,000 ms; echo time TE = 28 ms, 39 axial slices (3 mm thick aligned to the AC-PC plane with 0.6 mm slice gap), a 20 × 20 cm field-of-view, SENSE factor = 2, Flip angle = 77°. (The voxel size for resting-state scans was 3.5 mm^3^ rather than 3 mm^3^). To allow for T1 equilibrium effects, the first four volumes were acquired but not used in analyses. Foam padding was used to restrict head motion, and participants viewed the experimental stimuli on a screen through a mirror mounted on the head coil.

### fMRI data preprocessing

fMRI preprocessing was conducted using Analysis of Functional NeuroImages (AFNI) following steps described in previous work (Corriveau et al., 2024). Firstly, the first three volumes of each run were discarded to allow for magnetization equilibrium. Volumes with excessive head motion, defined as derivatives exceeding 0.25 mm in Euclidean norm, and volumes with outliers in more than 10% of voxels were censored from further analysis. Despiking was applied to reduce large signal spikes in the time series, followed by slice-time correction to adjust for timing differences across slices. Motion correction was performed; regressors included both demeaned and derivative terms of the six motion parameters, resulting in a 24-parameter motion model. Signals from subject-specific eroded regions of cerebrospinal fluid, white matter, and the whole brain were regressed out as nuisance variables. Functional images were then aligned to each subject’s high-resolution MPRAGE anatomical image and normalized to MNI space for group-level comparisons. Quality control criteria excluded runs with a maximum censored displacement exceeding 3 mm, an average censored motion greater than 0.15 mm, or more than 50% of TRs censored per run.

### Data exclusion

FMRI data were collected from 104 participants. Runs were excluded for missing fMRI and/or behavioral data, excessive head motion (>50% censored frames, average censored motion >.15mm, maximum displacement >3mm), and insufficient brain coverage (i.e., missing nodes). Participants were excluded from analyses comparing MA vs. placebo scans of an fMRI data type (rest, MID task, Doors task) if they did not have a usable run in both the MA and placebo conditions.

Exclusion criteria applied to resting-state data resulted in a final sample of *n*=82 in the MA condition, *n*=92 in the placebo condition, and *n*=76 in analyses comparing the two. The same criteria applied to MID task data resulted in a final sample of *n*=79 in the MA condition, *n*=85 in the placebo condition, and *n*=71 in neural analyses comparing the two. Of these, 69 participants are included in behavioral analyses comparing the two because two participants were missing gradCPT data.

The Doors task was collected in two fMRI runs. In the MA condition, exclusion criteria resulted in the removal of 19 run 1 scans and 21 run 2 scans. In the placebo condition, the same criteria resulted in the removal of 19 run 1 scans and 17 run 2 scans. For both the MA and placebo conditions, we averaged the run 1 and run 2 functional connectivity matrices if both runs were included, and only used the functional connectivity matrix for one run if only one run’s data was excluded. This resulted in 88 usable participants in the MA condition, 89 in the placebo condition, and 83 in both.

### Change in attention network strength due to methamphetamine

Functional network nodes were defined using the 268-node whole-brain Shen atlas (Shen et al., 2013). For our primary analysis, we included all 268 nodes and, for each subject, excluded nodes with missing data. Functional connectivity, defined as the Fisher z-transformed Pearson correlation between the time series of fMRI signals from each pair of atlas regions, was calculated for each fMRI run (resting-state, Doors, and MID) in each session (drug and placebo) independently.

To examine how functional connectivity signatures of sustained attention varied under drug and placebo conditions, we measured sustained attention network strength for each functional connectivity matrix. This was done using the high- and low-attention network masks available at https://github.com/monicadrosenberg/Rosenberg_PNAS2020, which together make up the established sustained attention CPM (Rosenberg et al., 2016a). Both the high- and low-attention network masks include 268 × 268 binary matrices, with a value of 1 indicating a functional connection, or edge, within the network. For each matrix, we applied the high- and low-attention masks separately and averaged the values in each network, producing high- and low-attention network strength measures for each functional connectome. Our analysis thus generated three pairs of variables of interest: six high- and low-attention network strength scores per participant across both MA and placebo conditions (two conditions × three run types [rest, MID task, Doors task]).

Previous findings have shown that sustained attention network strength reflects both individual differences and within-subject variability in sustained attention performance: Higher high-attention network strength and lower low-attention network strength are correlated with better sustained attention (Rosenberg et al., 2016a; Rosenberg et al., 2020). Further, individuals given methylphenidate had increased high-attention network strength and decreased low-attention network strength relative to unmedicated individuals (Rosenberg et al., 2016b). To conceptually replicate and extend previous findings showing modulation of sustained attention network strength by pharmacological conditions, we conducted paired t-tests comparing high- and low-attention network strength between MA and placebo conditions during rest. To investigate whether this effect held across different task contexts, we repeated these comparisons for both the MID task and Doors task.

### Selectivity of methamphetamine effects

To determine whether MA administration selectively modulates networks associated with sustained attention, we assessed if changes in the sustained attention networks were more pronounced than changes in other functional networks. This selectivity was evaluated in three ways.

First, we compared the effect of MA on two other connectome-based predictive models associated with arousal and valence (Ke et al., 2025), aiming to determine if MA’s effect was specific to attention-related networks. The arousal and valence networks were originally identified based on dynamic functional connectivity patterns that tracked self-reported arousal and positive/negative feelings during movie watching. The arousal network, which captures moment-to-moment fluctuations in emotional intensity, was found to generalize across individuals and situational contexts. Given that arousal is closely linked to attention, we expected that the arousal network would show a similar change in functional connectivity pattern under MA condition. In contrast, the valence network, which is primarily related with mood rather than momentary attention fluctuations, was not expected to exhibit significant changes. We computed the strength of both arousal and valence networks for the fMRI run and compared these between MA and placebo conditions to assess change in network strength.

Second, we examined MA’s effect on eight canonical resting-state networks: the medial frontal network, the frontoparietal network, the default mode network, the subcortical-cerebellar network, the motor network, the visual I network, the visual II network, and the visual association network (defined in Finn et al., 2015). This comparison aimed to identify any distinct MA-related effects on the sustained attention networks versus core resting-state networks. Each network strength was calculated as the mean functional connectivity strength between all pairs of nodes in that network and compared across conditions during rest, the MID task, and the Doors task.

Third, we employed Support Vector Classification (SVC) to identify connections within the functional connectome that significantly differed between MA and placebo conditions. SVC has been utilized in neuroimaging studies to classify different populations based on their functional connectivity patterns, such as distinguishing between patients with schizophrenia and healthy controls using resting-state fMRI data (Shen et al., 2010), and between individuals with major depressive disorder and healthy individuals (Craddock et al., 2009). In our study, we applied SVC to resting-state fMRI data to classify the two pharmacological conditions. Specifically, we used a leave-one-subject-out approach: for each fold, both MA and placebo scans from the same participant were held out together while the classifier was trained on the remaining participants’ scans. For participants with only one usable scan type, that scan was held out for testing. We then generated predictions for each of the held-out participant’s scan(s). This approach extends the application of SVC from inter-group comparisons to within-subject pharmacological effects, providing insights into how MA influences whole-brain functional connectivity. We consider this analysis a “specificity test” as it tests whether the functional connectivity differences (if any) that discriminate MA from placebo within subjects overlap with the functional connections in the high- and low-attention and arousal networks.

To assess the statistical significance of the classification accuracy, we performed a permutation test by randomly shuffling the MA and placebo labels of the test subject. By permuting the condition labels while keeping the functional connectivity matrices unchanged, we break any true association between the connectivity patterns and the pharmacological conditions. We then rerun the leave-one-out cross-validation SVC for each permutation to classify participants based on their functional connectivity matrices into the shuffled conditions. This process was repeated 1,000 times to build a distribution of classification accuracies that would be expected by chance under the null hypothesis of no fixed effect of MA on functional connectivity. By comparing the actual classification accuracy to this null distribution, we aimed to determine whether the observed classification accuracy was significantly higher than random chance.

To test whether the overlap between MA networks identified with SVC and the sustained attention and arousal networks was greater than expected by chance using the hypergeometric cumulative distribution function. This function assesses the probability of selecting up to 𝑥 shared edges out of 𝑀 total possible connections, given that one network includes 𝐾 edges and the other has 𝑛 edges, drawn without replacement. In our MATLAB code, this was computed as 𝑃 = 1 − ℎ𝑦𝑔𝑒𝑐𝑑𝑓 (𝑥, 𝑀, 𝐾, 𝑛). Here, 𝑥 is the number of overlapping edges, 𝐾 is the number of edges in the CPM network, 𝑛 is the number of edges in the propofol network, and 𝑀 is 35,778, which represents the total number of possible edges for a matrix of 268 brain regions.

### Behavioral prediction

We tested whether the sustained attention connectome-based predictive model (saCPM) generalized to predict out-of-scanner gradCPT performance. The saCPM, a linear model defined in Rosenberg et al. (2016a) that generates a predicted gradCPT *d’* score, consists of two parts: (1) high-attention and low-attention network masks and (2) a coefficient and error term. To generate predicted scores, we calculated high- and low-attention network strength during each rest and task run by averaging the functional connectivity values in each mask. For each subject and run type, we next subtracted low-attention network strength from high-attention network strength and input the resulting difference score into the linear model. We correlated the resulting predictions to actual gradCPT performance values obtained during the same drug state (MA or placebo). Given the study design, the MA and placebo functional connectivity data were obtained on different days than the MA and placebo gradCPT *d’* values.

## Results

### Methamphetamine modulates sustained attention performance

We compared participants’ changes in sustained attention performance in MA versus placebo conditions using the gradCPT. Within-subject gradCPT performance (*d’*) showed a modest improvement in the placebo condition compared to the orientation session (*t*(87)= 2.62, *p* = 0.01; **Figure 1)**, suggesting a placebo or practice effect. Performance in the MA condition was significantly higher than in the placebo condition (*t*(87) = 4.04, *p* < 0.001) and the orientation condition (*t*(87) = 6.20, *p* < 0.001; **Figure 1A)**, indicating better sustained attention task performance in the MA condition. We split all participants into high and low performer groups based on their gradCPT performance in the orientation session. Both groups showed improvement in performance in the MA vs. the placebo session (high performers: *t*(43) = 2.82, *p* = 0.007; low performers: *t*(43) = 2.95, *p* = 0.005; **Figure 1B**). We found similar results when we split participants into high and low performer groups using their gradCPT performance in the placebo session (**Figure S1**).

**Figure 1.**
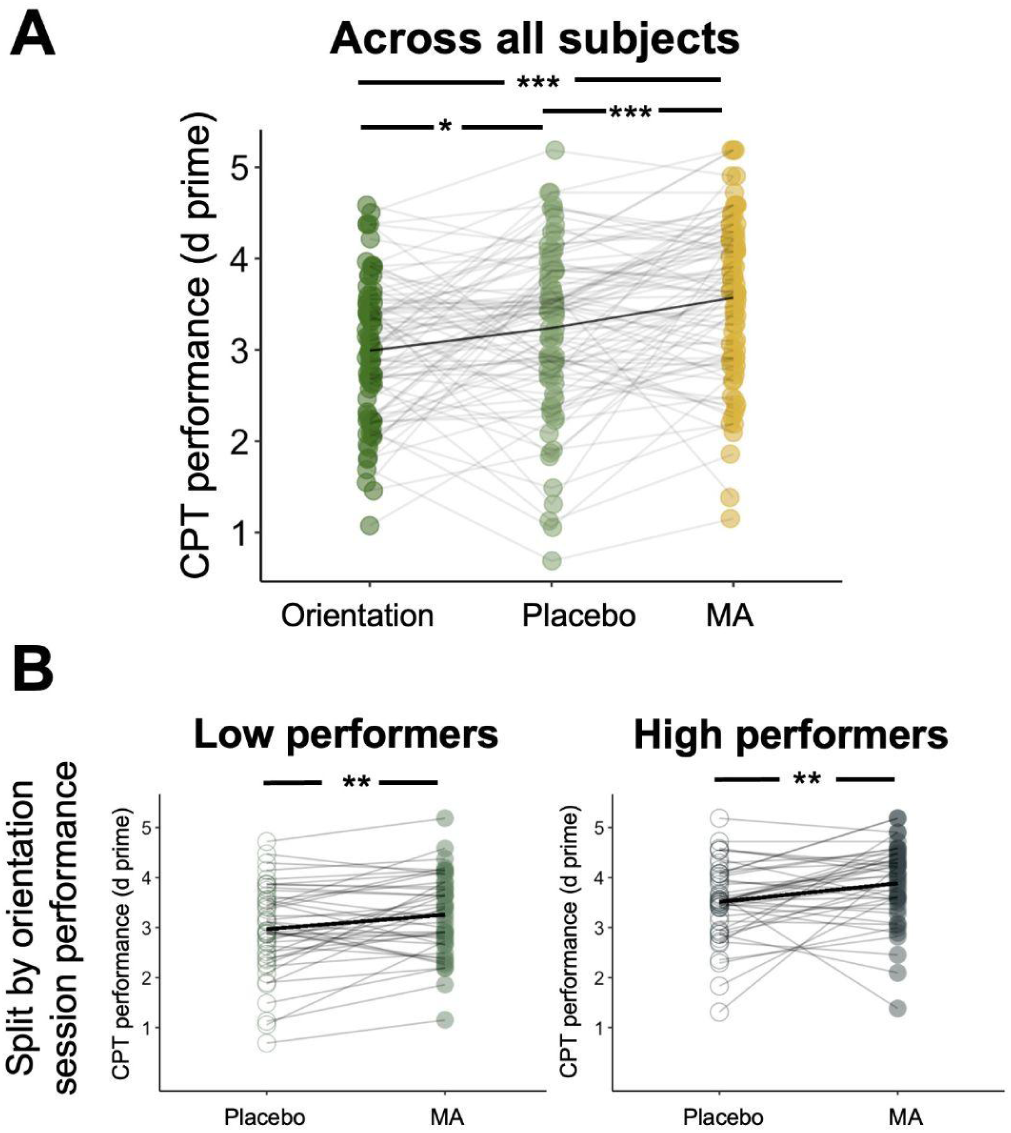
**A.** Gradual-onset continuous performance task (CPT) scores were higher under methamphetamine (MA) than either orientation or placebo sessions. **B.** Participants were split into high and low performer groups using a median split based on their CPT performance in the orientation session. Both low performers and high performers have improved CPT performance in the MA condition compared to the placebo condition.

We next explored whether the order of MA administration affected gradCPT performance. For participants who received MA in their first scan session, accuracy did not differ between MA and placebo conditions (*t*(52) = -0.27, *p* = 0.78). In contrast, for participants who received MA in the second session, accuracy during the MA session was significantly higher than during the first placebo session (*t*(53) = 2.76, *p* < 0.01). An MA-related performance improvement appeared only in participants who received MA after an initial placebo session.

### Methamphetamine increases functional connectivity signatures of sustained attention

We next examined the effects of MA on functional connectivity by comparing sustained attention network strengths between the MA and placebo conditions at rest and during task performance. As expected, MA increased high-attention network strength compared to the placebo condition at rest (*t*(75) = 8.41, *p* < 0.001; **Figure 2A**) and during both tasks (Doors task: *t*(82) = 10.35, *p* < 0.001; MID task: *t*(70) = 7.50, *p* < 0.001; **Supplementary Figures S2.A, S3.A**). In contrast, strength in the low-attention network was lower in the MA than the placebo condition at rest (*t*(75) = -9.54, *p* < 0.001; **Figure 2A**) and during the tasks (Doors task: *t*(82) = -9.75, *p* < 0.001; MID task: *t*(70) = -7.94, *p* < 0.001; **Supplementary Figures S2.A, S3.A**). Given that the MA condition showed increased high-attention network strength and reduced low-attention network strength, these effects are not likely due to global increases or decreases in functional connectivity.

**Figure 2.**
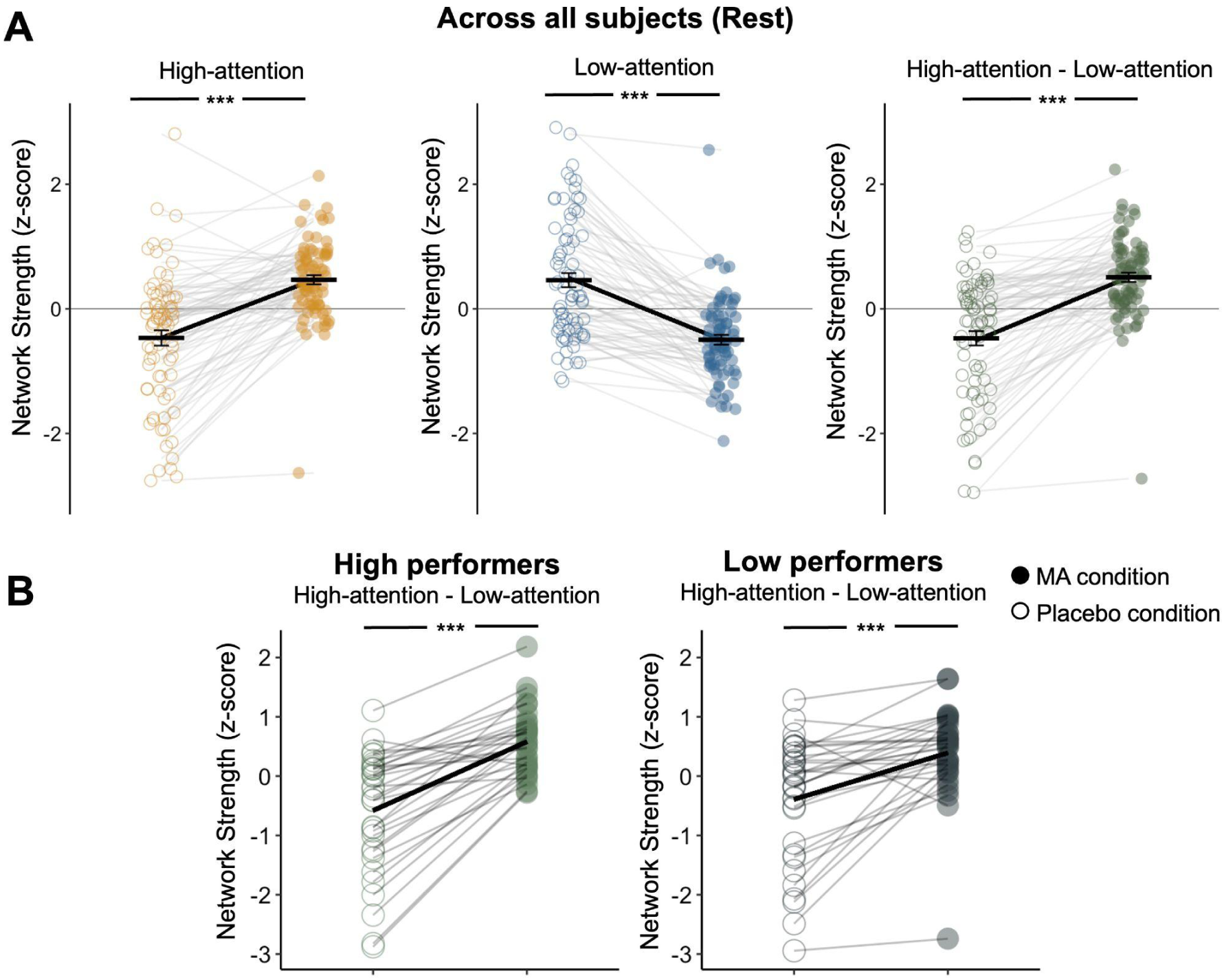
**A.** MA administration increased strength in the high-attention network and decreased strength in the low-attention network relative to control conditions. Network strength was calculated during the rest session, and was normalized within each graph for visualization. **B.** Effects of MA on network strength were similar regardless of baseline orientation session performance on the gradCPT. Dots represent individual participants, horizontal lines denote condition means, and error bars represent standard error of the mean.

We examined the effects of MA on high attention network strength in high and low performers at orientation. MA increased high-attention network strength and reduced low-attention network strength in both high performers (*t*(31) = 5.03, *p* < 0.001; **Figure 2B**) and low performers (*t*(31) = 8.52, *p* < 0.001; **Figure 2B**) during rest and during tasks (**Supplementary Figures S2.B, S3.B**).

### Methamphetamine selectively modulates sustained attention and arousal networks

#### Methamphetamine selectively modulates attention-related networks

Is MA’s effect on functional connectivity specific to networks associated with attention? To examine this, we compared MA’s impact on the sustained attention network to its effects on eight canonical resting-state networks, as well as valence and arousal networks identified in prior research (Ke et al., 2025). Given MA’s effect on attention task performance, we hypothesized that MA would modulate the arousal network but have little impact on the valence network, which is more associated with emotional processing rather than attention. Results showed that the difference in sustained attention network connectivity during rest (measured as mean strength of functional connections in the high-attention network connectivity minus mean strength of functional connections in the low-attention network) was significantly stronger in the MA condition than in the placebo condition (**Figure 3A**). A similar effect was observed for the arousal network (*t* (75)= 9.95, *p* < 0.001), but not for the valence network (*t* (74) = -1.05, *p* = 0.297), which is consistent with the idea that the arousal and sustained attention networks reflect overlapping constructs. Indeed, the sustained attention and arousal networks show overlapping features (number of edges in the sustained attention networks: 1387; number of edges in the arousal networks: 2014; number of overlapping edges in the expected direction [i.e., high-attention with high-arousal and low-attention with low-arousal]: 50; number in the unexpected direction: 13).

**Figure 3.**
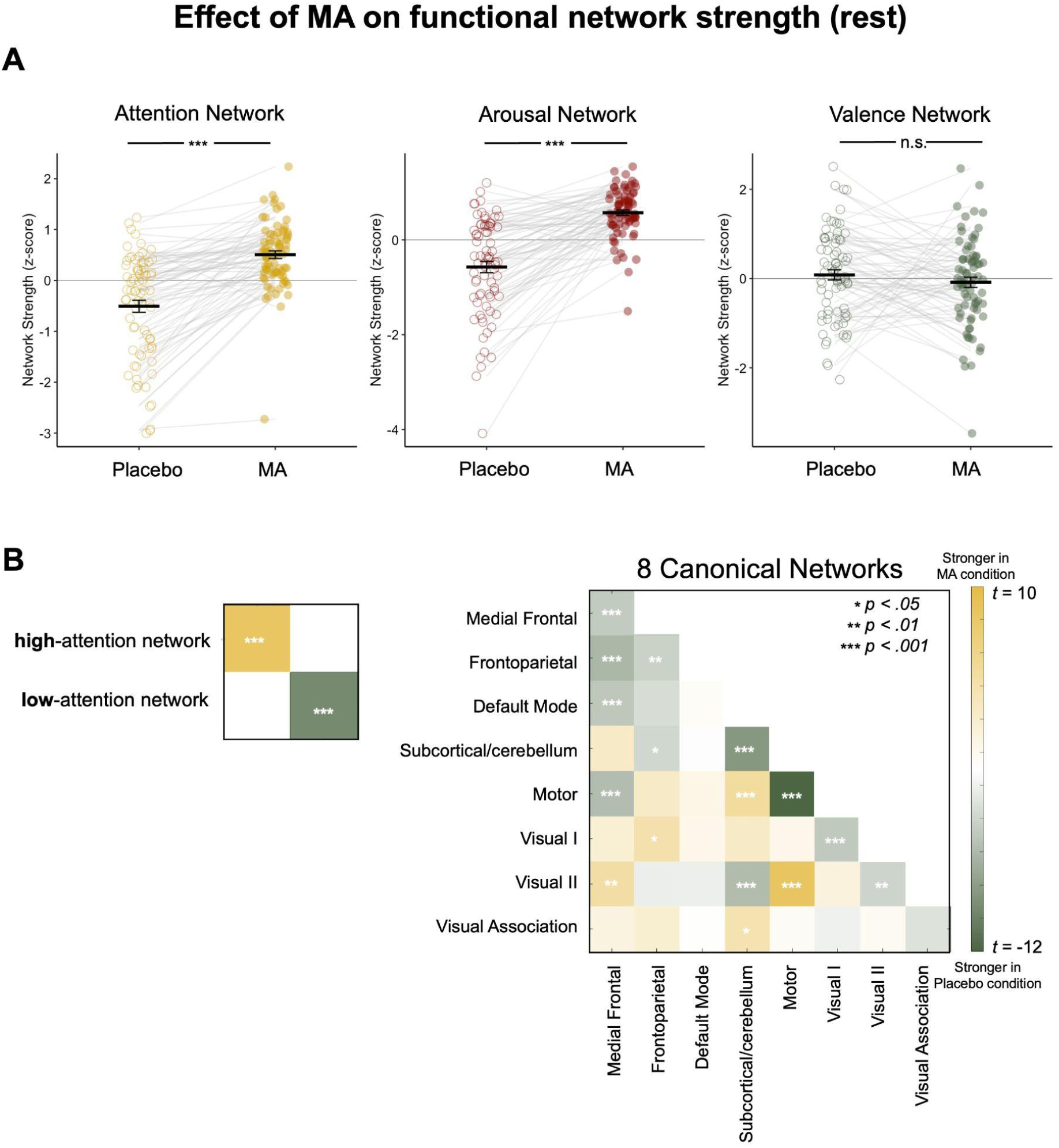
**A.** MA significantly modulates both the sustained attention and arousal networks compared to the placebo condition, whereas no significant effect is observed for the valence network. **B.** Statistically significant connectivity differences between MA and placebo conditions are indicated by stars. For each canonical network pair, we computed the actual connectivity difference (MA minus placebo) across subjects. For every pair of canonical networks, we computed each participant’s mean connectivity under the amphetamine (MA) and placebo (PL) conditions and then took the within-subject difference (MA – PL). We tested whether the group-mean difference for each pair deviated from zero with a two-tailed one-sample *t*-test. The resulting 64 *p*-values were Bonferroni-corrected (α = 0.05/64). Heat-map shows the corresponding *t*-statistics, with significance denoted by asterisks.

#### Methamphetamine modulates attention-related networks more than canonical resting-state networks

None of the 8 canonical resting-state networks showed significantly stronger within-network resting-state connectivity in the MA condition compared to placebo (**Figure 3B**; **Table 2**). Rather, within-network connectivity was weakened in the MA condition relative to placebo in a number of networks, including the default mode network (DMN) and visual network I and visual network II. MA significantly strengthened functional connectivity between the DMN and the subcortical cerebellar networks and between the DMN and visual network I. Interestingly, the only canonical network more affected by MA than the sustained attention networks was the motor network, which was stronger in the placebo condition (**Figure 3B**; **Table 2**). This aligns with prior observations of greater motor network connectivity in individuals with worse sustained attention function (Corriveau et al., 2024; O’Halloran et al., 2018).

**Table 1.** Demographic and drug use characteristics of the study participants.

| Demographic Categories | n (%) or mean (SD) |
| --- | --- |
| Sex (M/F) | 55/49 (53/47%) |
| Age (years) | 24.9 (± 4.1) |
| BMI | 23.5 (± 2.6) |
| Education (years) | 15.8 (± 1.6) |
| *Race/Ethnicity* |  |
| Asian | 17 (± 16%) |
| Black or African American | 5 (5%) |
| Hispanic | 21 (20%) |
| White | 54 (52%) |
| More than one race | 7 (7%) |
| *Drug Use (past month)* |  |
| Caffeinated drinks/day | 1.1 (± 1.0) |
| Alcoholic drinks/week | 3.6 (± 3.9) |
| Cannabis uses/month | 3.2 (± 5.6) |
| Daily nicotine users | 5 (5%) |
| *Lifetime stimulant use (times ever used)* | 1.8 (± 7.1) |

**Table 2.** Functional connectivity differences for the sustained attention networks (High, Low, and High–Low) and each of the eight canonical resting-state networks between MA and placebo conditions. Positive mean differences (*M*) indicate stronger connectivity under MA compared to placebo, and negative values indicate weaker connectivity under MA. *t*-scores and *p*-values come from one-sample *t*-tests against zero on the difference scores between conditions, and effect sizes represent Cohen’s *d*. All *p*-values are corrected for multiple comparisons using Bonferroni correction.

| Network | Mean MA–placebo difference | Effect size |
| --- | --- | --- |
| **Attentional Network** |  |  |
| High Attention network | $M = 0.040, t(75) = 8.415, p < .001$ | 0.97 |
| Low Attention Network | $M = -0.043, t(75) = -9.539, p < .001$ | -1.09 |
| Attention network (High - Low) | $M = 0.083, t(75) = 9.792, p < .001$ | 1.12 |
| Valence network (High - Low) | $M = -0.0001, t(75) = -1.049, p = .297$ | -0.12 |
| Arousal network (High - Low) | $M = 0.001, t(75) = 9.953, p < .001$ | 1.14 |
| **Canonical Networks** |  |  |
| Medial Frontal | $M = -0.028, t(75) = -4.746, p < .001$ | -0.54 |
| Frontoparietal | $M = -0.019, t(75) = -4.219, p = .004$ | -0.48 |
| Default Mode | $M = -0.003, t(75) = -0.515, p > .999$ | -0.06 |
| Subcortical-Cerebellum | $M = -0.021, t(75) = -8.318, p < .001$ | -0.95 |
| Motor | $M = -0.072, t(75) = -12.33, p < .001$ | -1.41 |
| Visual I | $M = -0.056, t(75) = -4.831, p < .001$ | -0.55 |
| Visual II | $M = -0.090, t(75) = -4.011, p = .009$ | -0.46 |
| Visual Association | $M = -0.031, t(75) = -3.071, p = .190$ | -0.35 |

### Methamphetamine modulates attention and arousal networks consistently across rest and task sessions

We previously observed that MA strengthened attention networks in both tasks and rest. We next asked whether the degree of this change was consistent across scan types. In both sustained attention and arousal networks, we found strong correlations in MA-induced network changes between task and rest. Specifically, changes in the sustained attention and arousal networks were significantly correlated between the rest session and the Doors task session (attention: *r* = 0.544, *p* < 0.001; arousal: *r* = 0.481, *p* < 0.001), between rest session and the MID task session (attention: *r* = 0.468, *p* < 0.001; arousal: *r* = 0.435, *p* = 0.001), and between the Doors and MID task sessions (attention: *r* = 0.460, *p* < 0.001; arousal: *r* = 0.438, *p* < 0.001). This suggests that MA’s effects on attention-related networks are consistent across different conditions or contexts, indicating a stable influence of the drug on functional networks whether the brain is at rest or engaged in task-based activity.

In contrast, MA’s effect on the valence network varied across sessions. While there was a significant correlation in MA-induced changes between the Doors and MID task sessions (*r* = 0.494, *p* < 0.001), no significant correlation was observed between the rest session and the Doors task session (*r* = -0.039, *p* = 0.777) or between the rest session and the MID task session (*r* = 0.072, *p* = 0.605). This suggests that MA’s effect on the valence network may be task-dependent or simply noise, showing consistency between active tasks but not between rest and task conditions.

### Methamphetamine network anatomy

Using a support vector classifier (SVC) with a linear kernel, we observed a leave-one-out cross-validation classification accuracy of 83.91% in distinguishing between MA and placebo conditions based on the resting-state functional connections. To assess the statistical significance of this result, we conducted a permutation test by randomly shuffling the drug and placebo labels and rerunning the classification to establish a chance-level distribution. The actual classification accuracy was significantly higher than chance (*p* < 0.001; **Figure S4**).

We identified the most influential functional connections, or edges, in the classification by examining the weight coefficients of the SVC. We selected the top 2.5% of both positive and negative average weights across all leave-one-out cross-validation prediction runs. Specifically, edges with average weights exceeding 0.0020 or below -0.0023 were designated as significant, resulting in 135 positive edges and 168 negative edges from the total set of possible connections. The analyses revealed that connections between the motor cortex and cerebellum, limbic regions, basal ganglia, and fronto-parietal networks were particularly influential for the classification. Additionally, significant connections emerged between the medial frontal cortex and both the motor and visual cortices (see **Figure S4, Figure S5**).

Notably, the positive and negative edges identified from the SVC (MA network) did not significantly overlap with the high-attention and low-attention networks identified by Rosenberg et al. (2016a) as predictors of sustained attention (**Table 3**). However, the MA network significantly overlaps with the edges positively and negatively related to arousal in the arousal network in Ke et al. (2025) (**Table 3**). Given that the arousal network is linked to the intensity and activation of emotional states, this significant overlap may suggest that MA’s influence on functional brain connectivity aligns more closely with neural patterns associated with arousal rather than sustained attention. This finding implies that MA may modulate large-scale brain networks in a way that parallels the experience of heightened physiological and emotional activation.

**Table 3.**
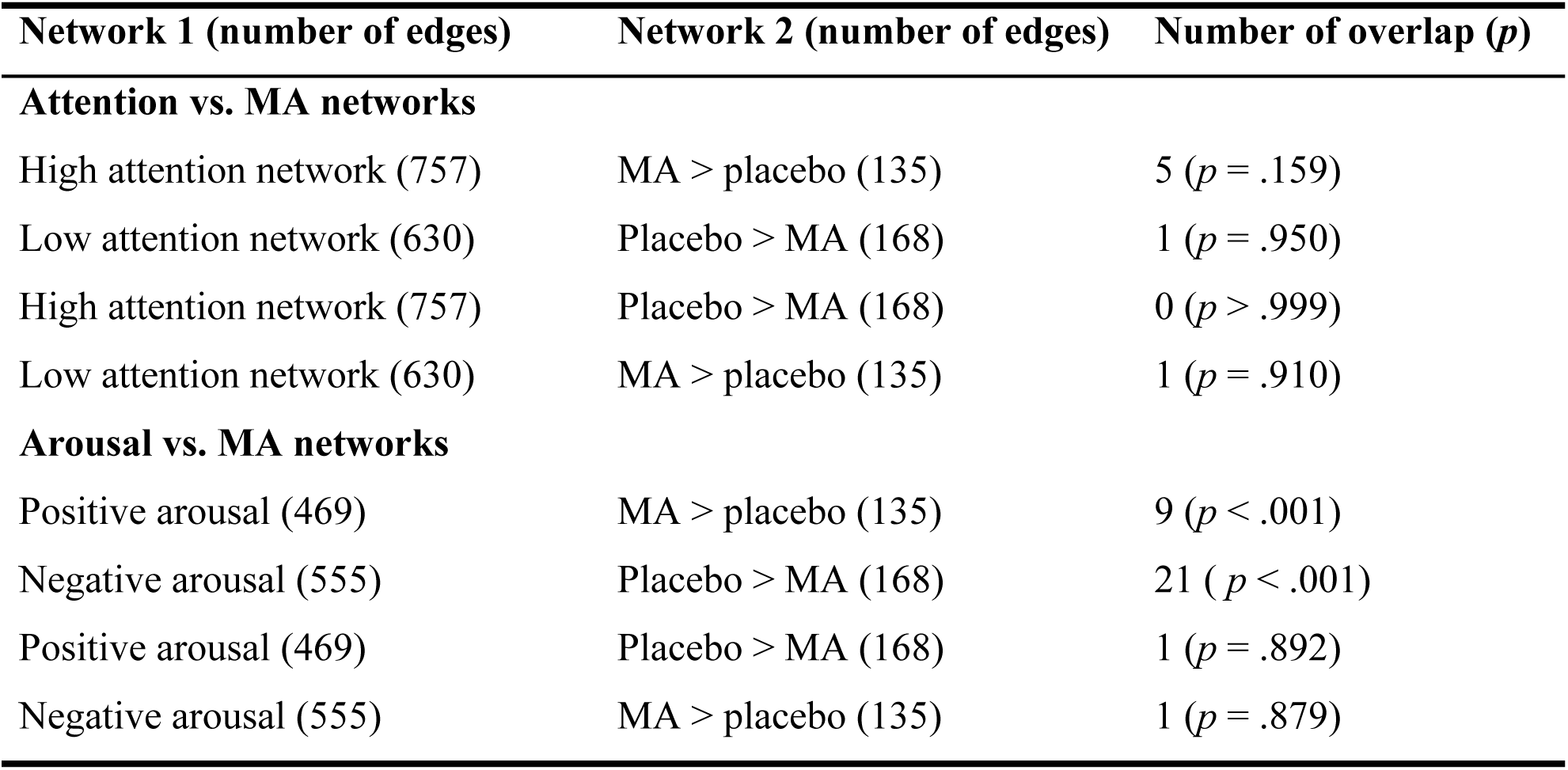
Overlap between the sustained attention and arousal networks and the classification edges derived from the SVC (MA(+)-weighted [MA > placebo] vs. MA(-)-weighted [placebo > MA]).

| Network 1 (number of edges) | Network 2 (number of edges) | Number of overlap (*p*) |
| --- | --- | --- |
| **Attention vs. MA networks** |  |  |
| High attention network (757) | MA > placebo (135) | 5 (*p* = .159) |
| Low attention network (630) | Placebo > MA (168) | 1 (*p* = .950) |
| High attention network (757) | Placebo > MA (168) | 0 (*p* > .999) |
| Low attention network (630) | MA > placebo (135) | 1 (*p* = .910) |
| **Arousal vs. MA networks** |  |  |
| Positive arousal (469) | MA > placebo (135) | 9 (*p* < .001) |
| Negative arousal (555) | Placebo > MA (168) | 21 (*p* < .001) |
| Positive arousal (469) | Placebo > MA (168) | 1 (*p* = .892) |
| Negative arousal (555) | MA > placebo (135) | 1 (*p* = .879) |

### Behavioral prediction

#### Sustained attention network strength does not predict CPT performance

We found that saCPM predictions did not correlate with gradCPT *d’* when generated from resting-state connectivity (MA: *r* = -0.109, *p* = .329; Placebo: *r* = -0.036, *p* = .735), Doors task connectivity (MA: *r* = -0.097, *p* = .367; Placebo: *r* = -0.016, *p* = .884), or MID task connectivity (MA: *r* = 0.036, *p* = .753; Placebo: *r* = 0.087, *p* = .432). This finding suggests that while the saCPM has been extensively validated in prior work as a strong predictor of sustained attention (e.g., Rosenberg et al., 2016a; Rosenberg et al., 2020; Kardan et al., 2022), it does not predict gradCPT performance under MA or placebo in the current sample. This may be due to the fact that the fMRI scans and the gradCPT were performed on two different days (but under the same drug condition), which may have introduced additional variability in the observed effects.

#### Change in sustained attention network strength does not predict change in CPT performance

Similarly, changes in the saCPM’s high- and low-attention networks induced by MA administration, measured as the difference in functional connectivity between the MA and placebo conditions, did not predict corresponding changes in gradCPT performance using resting-state connectivity (*r* = 0.167, *p* = .149, N = 76), Doors task connectivity (*r* = -0.016, *p* = .888, N = 83), or MID task connectivity (*r* = 0.080, *p* = .514, N = 69). Therefore, while we observed significant increases in both sustained attention performance and sustained attention network strength under the MA condition, these changes were unrelated to one another. This is perhaps unsurprising due to previously reported results that (1) network strength in the saCPM did not predict gradCPT performance in the current sample and (2) saCPM and MA networks were largely non-overlapping.

#### MA’s modulation of the sustained attention networks differs between high and low performers

Although sustained attention network strength did not predict gradCPT performance, we next asked whether these networks may still capture individual differences in attention-related performance. One way to explore this is to test whether MA-induced changes in attention network strength vary depending on an individual’s initial gradCPT performance. We hypothesized that participants who initially performed well in the gradCPT without MA (high performers) might exhibit a smaller benefit due to a ceiling effect, whereas those who performed less well without drug (low performers) could show a greater increase if they had more room to improve. To address this, we categorized participants into high or low performers based on a median split of their gradCPT scores from the orientation session. During rest, both high performers (*t*(31) = 5.03, *p* < 0.001) and low performers (*t*(31) = 8.52, *p* < 0.001) showed significant increases in high-attention network strength and decreases in low-attention network strength in the MA condition compared to the placebo condition. This pattern was consistent during both task sessions (**Figures S2, S3**). Finally, to examine whether the magnitude of MA-induced changes in attention network strength depended on attentional performance under placebo, we tested the correlation between participants’ change in network strength and their gradCPT performance under placebo condition. However, no significant correlation was found using rest session (*r* = -0.059, *p* = .611, *N* = 76), or Doors session (*r* = -0.051, *p* = .650, *N* = 83) network strength. Although there was a relationship between placebo gradCPT d’ and MID session attention network strength change (*r* = -0.278, *p* = .021, *N* = 69), the lack of consistency across run types suggesting that MA’s enhancement on attention network strength is not robustly driven by initial performance levels.

## Discussion

In this study, we built on earlier evidence that another stimulant drug, methylphenidate, modulates attention-related functional brain networks (Rosenberg et al., 2016b). While the previous study used a between-subject design, where participants were split into two groups receiving either methylphenidate or no drug, the current study employed a within-subject, placebo-controlled design. In addition, the previous study used methylphenidate whereas the present study used MA to investigate sustained attention networks. With each participant serving as their own control, the within-subject design reduces noise from subject-to-subject variability, thereby providing a more direct comparison of the two conditions with higher statistical power. Rosenberg et al. (2016b) demonstrated that healthy individuals given a single dose of methylphenidate before fMRI showed whole-brain connectivity patterns indicative of better sustained attention—higher high-attention and lower low-attention network strength—than controls. Our within-subject design yielded the same pattern of attention-related network modulation. MA’s effect was most significant in the functional networks previously implicated in attentional processing, aligning with prior work on methylphenidate (Rosenberg et al., 2016b).

Beyond attention networks, our findings demonstrate that MA also modulated arousal-related functional connectivity. Specifically, under MA administration, high-attention and high-arousal network strength increased, while low-attention and low-arousal network strength decreased relative to the placebo condition. These changes were observed at rest and during two task sessions, suggesting that MA exerts a consistent influence on attention- and arousal-related brain networks across different cognitive states. The fact that arousal network connectivity changes paralleled those seen in sustained attention networks suggests that the two systems may be closely linked, with arousal modulation potentially serving as a mechanism through which MA enhances attentional performance.

However, while moderate increases in arousal may facilitate attentional and cognitive performance, excessive arousal could impair performance by pushing individuals into a hyperarousal state. According to the Yerkes-Dodson law, optimal task performance occurs at intermediate levels of arousal, whereas both under- and over-arousal can lead to decrements in performance (Khazaei et al., 2021). The observed MA-induced increase in arousal network strength raises the possibility that its effects on performance may not be linear. Individuals who were already near their optimal arousal state under placebo may have seen less benefit, or even potential impairment, if MA pushed them into a state of excessive arousal. Future research could explore this possibility by testing for a quadratic (inverted-U) relationship between arousal network strength and behavioral performance, which could clarify whether MA’s effects on attention are mediated by an optimal balance between increased arousal and cognitive control. Moreover, behavior on the gradCPT also improved after MA administration in individuals with good and poor initial performance. This is partially consistent with previous study on MA-dependent individuals, which demonstrated that MA can selectively enhance attention or information processing speed and working memory in those who exhibit poorer cognitive performance at baseline (Mahoney et al., 2011). Taken together, the current study suggests that MA selectively enhances resting-state and task-related connectivity in attention and arousal networks and affects sustained attentional performance.

Despite robust condition-level differences in attention-related network strength and behavior, our attempt to use the same connectome-based predictive model (saCPM) to predict performance on the continuous performance task did not yield significant results. Earlier work has shown that connectivity in these networks can predict within-subject attentional state changes (Rosenberg et al., 2020; Kardan et al., 2022). One possibility for the current null finding is that conducting the gradCPT and the fMRI scan sessions on separate days might have introduced variability in the drug’s effects, especially considering factors such as participant expectations across repeated administrations. Individual differences in how participants responded to a second dose of MA on a subsequent day may have affected the functional connectivity–behavior relationships in the saCPM. These findings motivate future work minimizing the timing of behavioral measurements relative to scanning sessions.

Our results shed light on how MA modulates large-scale functional connectivity related to sustained attention and arousal. A support vector classifier was able to distinguish between MA and placebo conditions with 83.91% accuracy, suggesting robust changes in functional connectivity patterns. Notably, we found that within-motor network functional connectivity was stronger in the placebo than the MA condition and thus potentially associated with weaker sustained attention. On the other hand, functional connectivity between the motor and cerebellar network and between motor and visual II network was stronger in the MA condition. These findings replicate and extend previous research demonstrating similar results suggesting heightened within motor network connectivity is linked to poorer sustained attention (Corriveau et al., 2024; O’Halloran et al., 2018). Interestingly, while previous results implicated stronger within-motor connections to worse sustained attention during tasks involving motor responses, the current findings suggest that a similar relationship can be observed at rest when no responses are necessary.

Despite MA’s enhancement of sustained attention networks and potential improvements in sustained attention-related performance at therapeutic doses, MA could impair cognitive functions that require visual scanning or the ability to filter irrelevant information (Silber et al., 2006). Chronic use in individuals with MA use disorder has further been linked to poorer short-term memory and executive functioning (Nestor et al., 2023) as well as mood disturbances and disrupted connectivity within key affective networks (Jiang et al., 2021). While MA’s heightened potency compared to amphetamine and negative side-effects are well-documented (Ipser et al., 2018; Silber et al., 2006), our data parallel findings from safer stimulants, like methylphenidate, in showing that MA can modulate sustained attention networks. Together, these findings suggest that improved sustained attentional performance under stimulants may arise from shared neurochemical pathways; however, the serious risks of addiction and other negative impacts highlight the need for caution when considering MA, even in low doses, as a potential therapeutic agent to enhance sustained attention.

In comparing the results of previous studies with methylphenidate to the present findings with MA, an important limitation is that both studies used only a single dose. Comparisons across drugs are difficult without a full dose-response function or without a reference dose using a measure on which both drugs are matched. However, we note that the subjective, behavioral and physiological effects of oral doses of 20 mg methamphetamine and 45 mg methylphenidate in humans are very similar (Martin et al., 1971; Heishman & Henningfield, 1991; Arkell et al., 2022). Thus, it is likely that the drugs also have comparable effects on neural function at these doses.

Beyond characterizing the behavioral and network-level effects of MA, our findings underscore the importance of testing the sensitivity and specificity of brain-based biomarkers of behavior—such as the connectome-based predictive models of sustained attention, arousal, and valence used here—across datasets and experimental contexts. That MA modulated the sustained attention and arousal CPMs, but not the valence CPM, supports our *a priori* prediction that MA would selectively affect attention- and arousal-related processes. Conversely, a finding that attention and arousal network strength remained unchanged or the valence network had been modulated would have motivated a reevaluation of the robustness, generalizability, and/or process-specificity of these models. Pharmacological manipulations like psychostimulants thus offer a powerful and underutilized approach for evaluating the validity of brain-based biomarkers by providing causal tests of their sensitivity and specificity.

In sum, we found that methamphetamine robustly modulates attention- and arousal-related functional connectivity networks, paralleling prior work with other psychostimulants (Rosenberg et al., 2016b). Using a support vector classifier, we reliably distinguished between methamphetamine and placebo conditions, reflecting systematic changes in network connectivity and showing that heightened motor network connectivity indicated worse sustained attention, which is consistent with prior research. Together, these findings demonstrate how psychostimulants can shape large-scale functional brain networks and how pharmacological manipulations can offer powerful opportunities for testing the validity of brain-based predictive models.

## Supplementary Data

**Figure S1.**
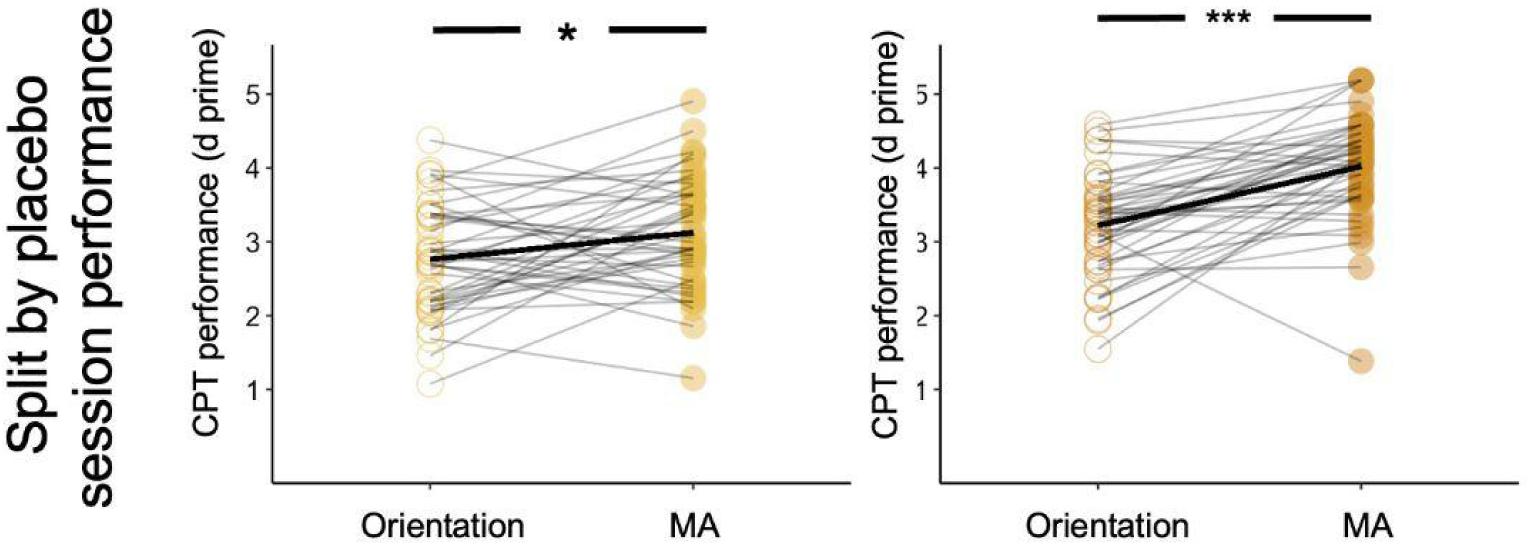
Participants were split into high and low performer groups using a median split based on their CPT performance under the placebo session. Both low performers and high performers have improved CPT performance in the MA condition compared to the orientation session.

**Figure S2.**
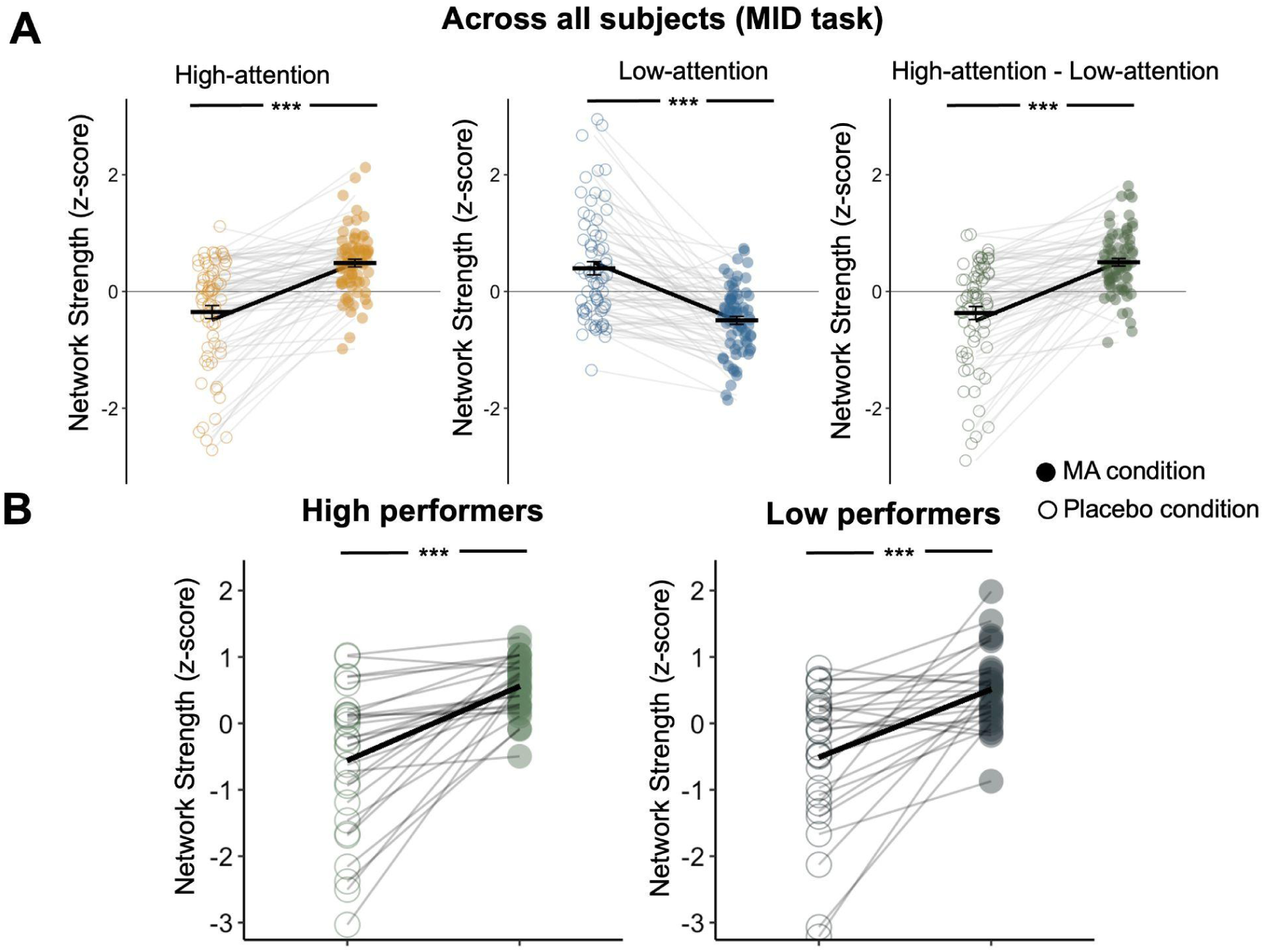
MA administration increased strength in the high-attention network and decreased strength in the low-attention network relative to control conditions. Network strength was calculated during the MID task, and was normalized within each graph for visualization. **B.** Effects of MA on network strength were similar regardless of orientation session performance on the gradCPT. Dots represent individual participants, horizontal lines denote condition means, and error bars represent standard error of the mean.

**Figure S3.**
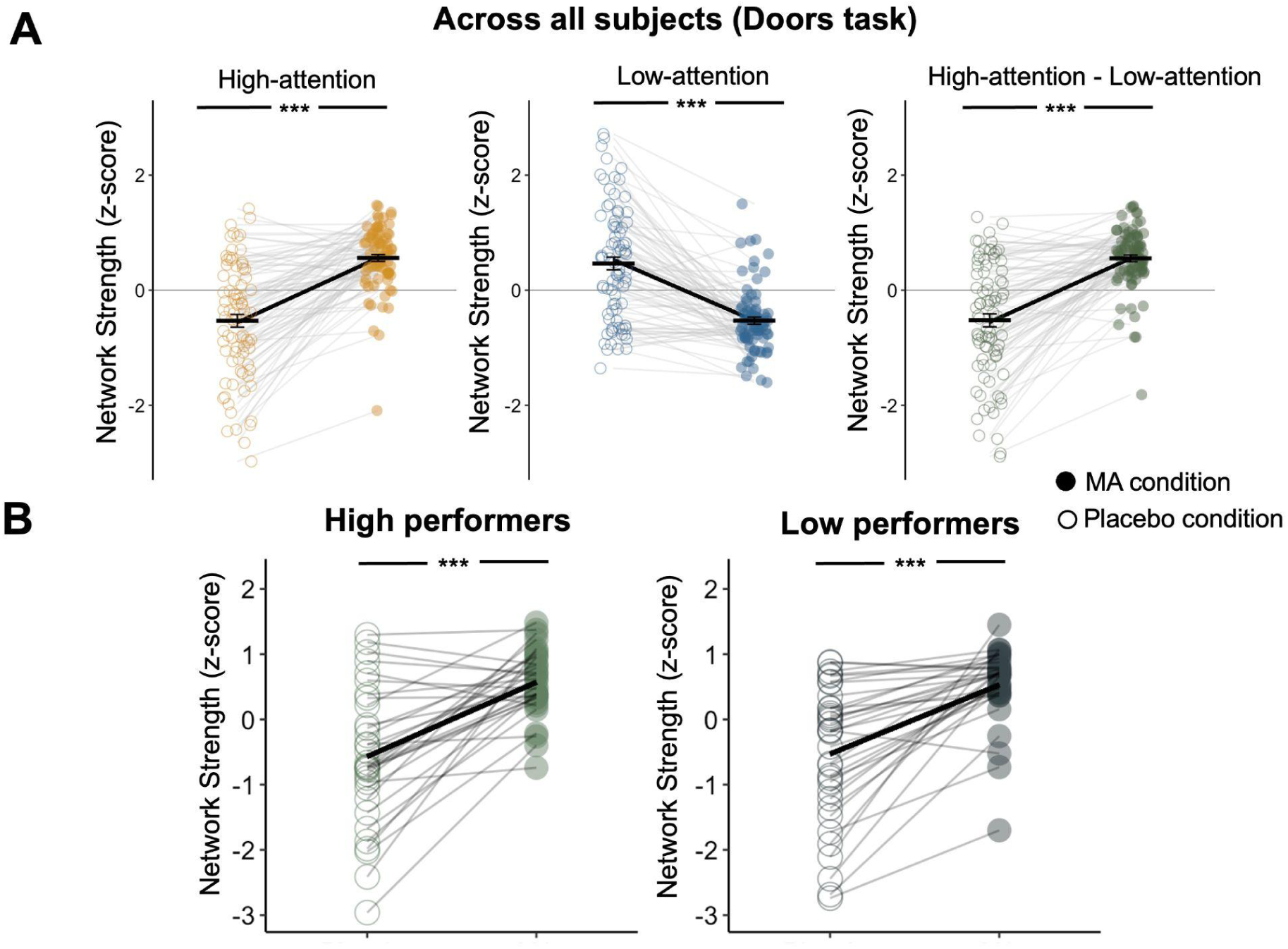
MA administration increased strength in the high-attention network and decreased strength in the low-attention network relative to control conditions. Network strength was calculated during the Doors task, and was normalized within each graph for visualization. **B.** Effects of MA on network strength were similar regardless of orientation session performance on the gradCPT. Dots represent individual participants, horizontal lines denote condition means, and error bars represent standard error of the mean.

**Figure S4.**
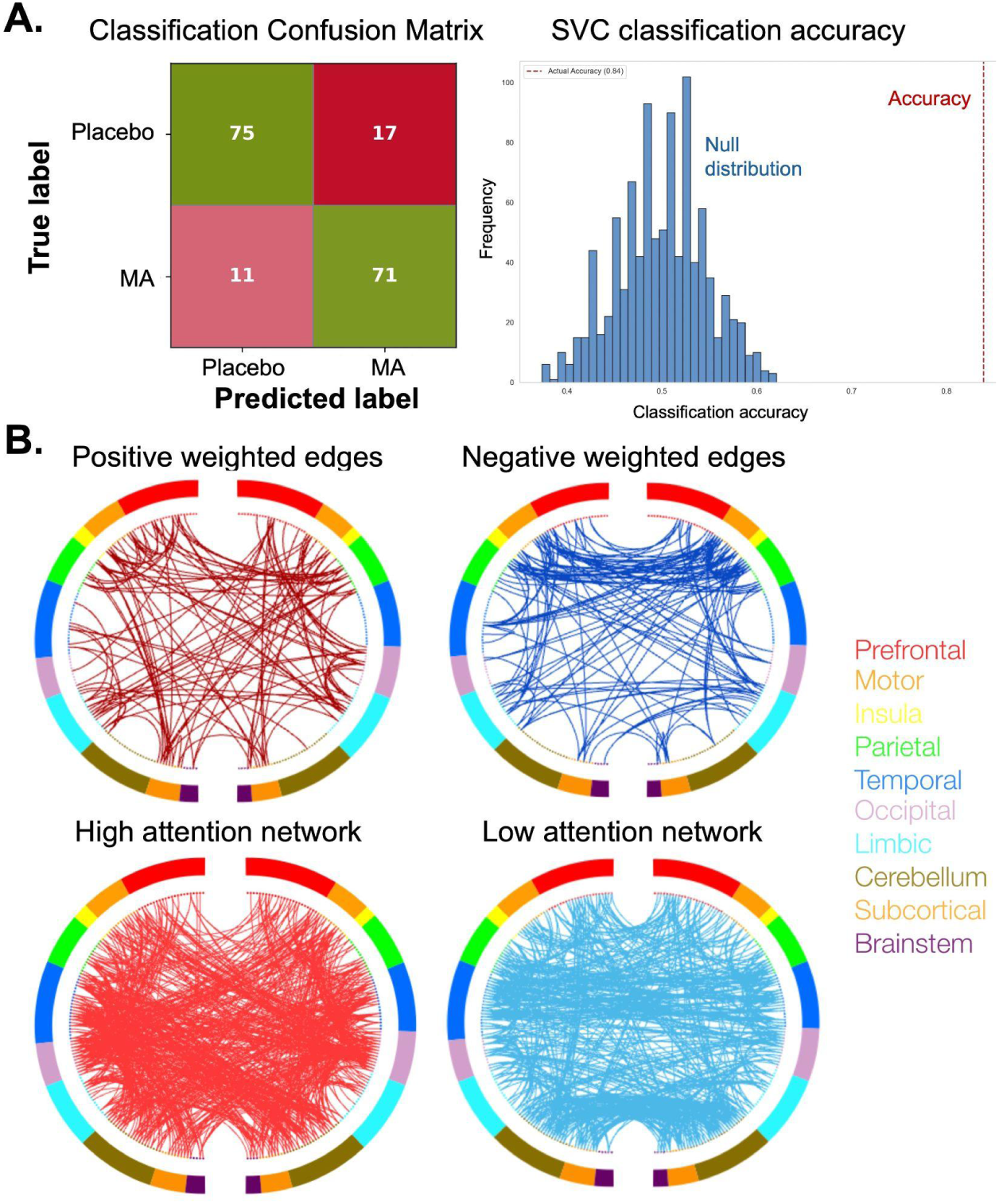
**A.** The support vector classifier distinguishes subjects in MA versus placebo conditions with an accuracy of 83.91%. **B.** The first row of circle plots shows the top 2.5% most positive and most negative weighted functional connectivity edges of the support vector classifier (SVC) in distinguishing between MA and placebo conditions. Networks were defined using functional connectivity patterns during the resting-state scan; similar results were observed for the Monetary Incentive Delay (MID) task and Doors task (Figure 5.1, 5.2). The second row of circle plots shows the high-attention and low-attention networks, defined to predict performance on an attention task in an independent group of participants (Rosenberg et al., 2016b). Nodes are grouped by macroscale regions, including the prefrontal cortex, motor cortex, insula, parietal cortex, temporal cortex, occipital cortex, limbic lobe (including the cingulate cortex, amygdala, and hippocampus), cerebellum, subcortex (thalamus and striatum), and brainstem. The right half of the circle represents the right hemisphere of the brain.

**Figure S5.**
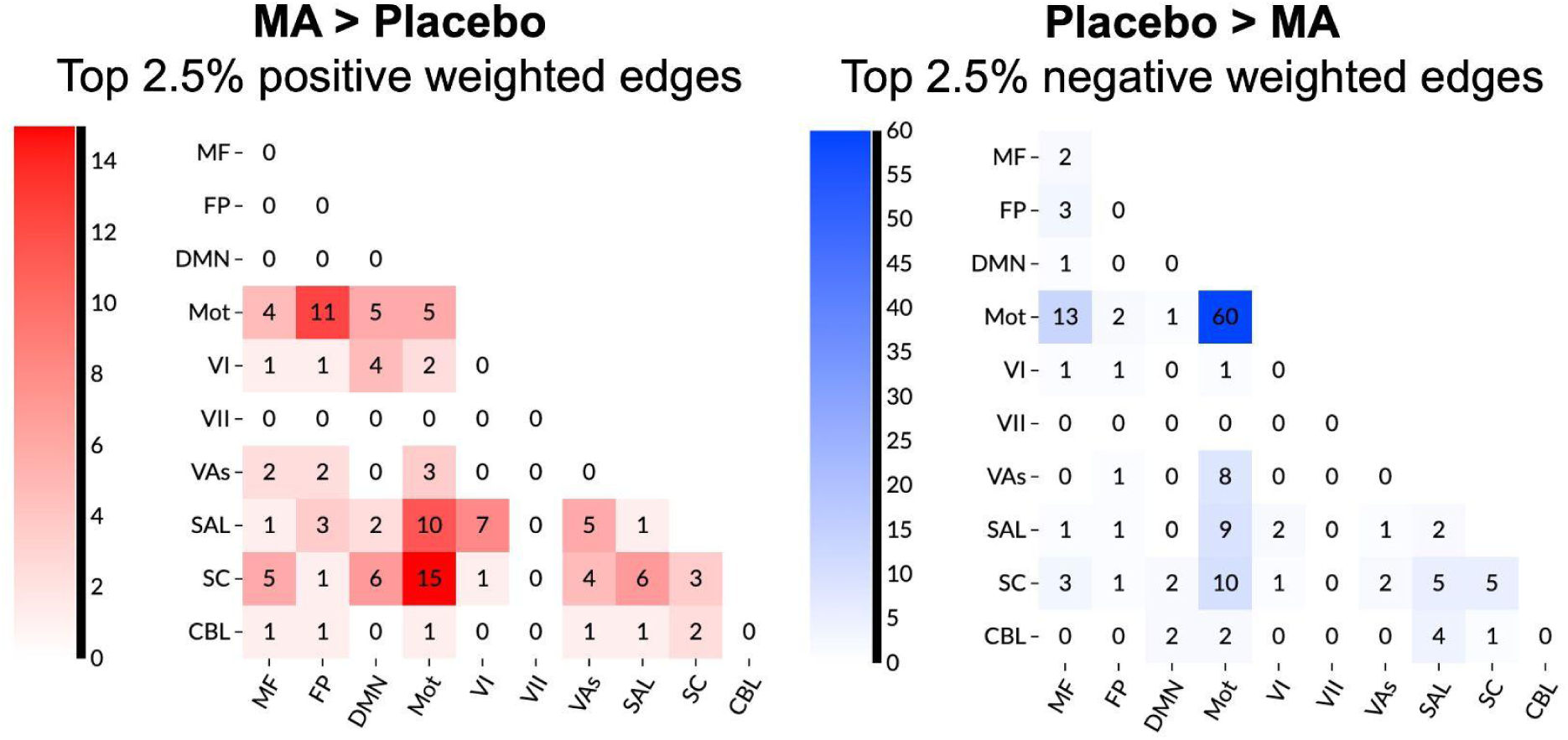
Top 2.5% positive and negative weighted edges in the MA network. Connections between the motor cortex and cerebellum, limbic regions, basal ganglia, and fronto-parietal networks were particularly influential for the classification.

## References

Ahmadlou, M., Ahmadi, K., Rezazade, M., & Azad-Marzabadi, E. (2013). Global organization of functional brain connectivity in methamphetamine abusers. Clinical Neurophysiology, 124(6), 1122–1131. 10.1016/j.clinph.2012.12.003

Arkell, T. R., Bradshaw, K., Downey, L. A., Hayley, A. C. (2022). Acute effects of amphetamine and related psychostimulants on impulsivity: a systematic review of clinical trials. Addiction Biology, 27: e13128. 10.1111/adb.13128

Avelar, A. J., Juliano, S. A., & Garris, P. A. (2013). Amphetamine augments vesicular dopamine release in the dorsal and ventral striatum through different mechanisms. Journal of Neurochemistry, 125(3), 373–385. 10.1111/jnc.12197

Berridge, C. W., Devilbiss, D. M., Andrzejewski, M. E., Arnsten, A. F. T., Kelley, A. E., Schmeichel, B., Hamilton, C., & Spencer, R. C. (2006). Methylphenidate Preferentially Increases Catecholamine Neurotransmission within the Prefrontal Cortex at Low Doses that Enhance Cognitive Function. Biological Psychiatry, 60(10), 1111–1120. 10.1016/j.biopsych.2006.04.022

Carlson, J. M., Foti, D., Mujica-Parodi, L. R., Harmon-Jones, E., & Hajcak, G. (2011). Ventral striatal and medial prefrontal BOLD activation is correlated with reward-related electrocortical activity: A combined ERP and fMRI study. NeuroImage, 57(4), 1608–1616. 10.1016/j.neuroimage.2011.05.037

Chamberlain, T. A., & Rosenberg, M. D. (2022). Propofol selectively modulates functional connectivity signatures of sustained attention during rest and narrative listening. Cerebral Cortex, 32(23), 5362–5375. 10.1093/cercor/bhac020

Contini, V., Rovaris, D. L., Victor, M. M., Grevet, E. H., Rohde, L. A., & Bau, C. H. D. (2013). Pharmacogenetics of response to methylphenidate in adult patients with Attention-Deficit/Hyperactivity Disorder (ADHD): A systematic review. European Neuropsychopharmacology, 23(6), 555–560. 10.1016/j.euroneuro.2012.05.006

Corriveau, A., Ke, J., Terashima, H., Kondo, H. M., & Rosenberg, M. D. (2024). Functional brain networks predicting sustained attention are not specific to perceptual modality. Network Neuroscience.

Covey, D. P., Juliano, S. A., & Garris, P. A. (2013). Amphetamine Elicits Opposing Actions on Readily Releasable and Reserve Pools for Dopamine. PLoS ONE, 8(5), e60763. 10.1371/journal.pone.0060763

Craddock, R. C., Holtzheimer, P. E., Hu, X. P., & Mayberg, H. S. (2009). Disease state prediction from resting state functional connectivity. Magnetic Resonance in Medicine, 62(6), 1619–1628. 10.1002/mrm.22159

Crane, N. A., Molla, H., & De Wit, H. (2023). Methamphetamine alters nucleus accumbens neural activation to monetary loss in healthy young adults. Psychopharmacology, 240(9), 1891–1900. 10.1007/s00213-023-06398-4

Devous, M. D., Trivedi, M. H., & Rush, A. J. (2000). Regional Cerebral Blood Flow Response to Oral Amphetamine Challenge in Healthy Volunteers. J Nucl Med.

Esterman, M., Noonan, S. K., Rosenberg, M., & DeGutis, J. (2013). In the zone or zoning out? Tracking behavioral and neural fluctuations during sustained attention. Cerebral cortex, 23(11), 2712–2723.

Esterman, M., & Rothlein, D. (2019). Models of sustained attention. Current opinion in psychology, 29, 174–180.

Faraone, S. V. (2018). The pharmacology of amphetamine and methylphenidate: Relevance to the neurobiology of attention-deficit/hyperactivity disorder and other psychiatric comorbidities. Neuroscience & Biobehavioral Reviews, 87, 255–270. 10.1016/j.neubiorev.2018.02.001

Faraone, S. V., Asherson, P., Banaschewski, T., Biederman, J., Buitelaar, J. K., Ramos-Quiroga, J. A., Rohde, L. A., Sonuga-Barke, E. J. S., Tannock, R., & Franke, B. (2015). Attention-deficit/hyperactivity disorder. Nature Reviews Disease Primers, 1(1), 15020. 10.1038/nrdp.2015.20

Garrett, D. D., Nagel, I. E., Preuschhof, C., Burzynska, A. Z., Marchner, J., Wiegert, S., Jungehülsing, G. J., Nyberg, L., Villringer, A., Li, S.-C., Heekeren, H. R., Bäckman, L., & Lindenberger, U. (2015). Amphetamine modulates brain signal variability and working memory in younger and older adults. Proceedings of the National Academy of Sciences, 112(24), 7593–7598. 10.1073/pnas.1504090112

Finn, E. S., Shen, X., Scheinost, D., Rosenberg, M. D., Huang, J., Chun, M. M., Papademetris, X., & Constable, R. T. (2015). Functional connectome fingerprinting: Identifying individuals using patterns of brain connectivity. Nature Neuroscience, 18(11), 1664–1671. 10.1038/nn.4135

Heishman, S. J. & Henningfield, J. E. (1991). Discriminative stimulus effects of d-amphetamine, methylphenidate, and diazepam in humans. Psychopharmacology, 103: 436–442. 10.1007/BF02244241

Ipser, J. C., Uhlmann, A., Taylor, P., Harvey, B. H., Wilson, D., & Stein, D. J. (2018). Distinct intrinsic functional brain network abnormalities in methamphetamine-dependent patients with and without a history of psychosis. Addiction Biology, 23(1), 347–358. 10.1111/adb.12478

Jiang, P., Sun, J., Zhou, X., Lu, L., Li, L., Huang, X., Li, J., Kendrick, K., & Gong, Q. (2021). Functional connectivity abnormalities underlying mood disturbances in male abstinent methamphetamine abusers. Human Brain Mapping, 42(11), 3366–3378. 10.1002/hbm.25439

Kardan, O., Stier, A. J., Cardenas-Iniguez, C., Schertz, K. E., Pruin, J. C., Deng, Y., … & Rosenberg, M. D. (2022). Differences in the functional brain architecture of sustained attention and working memory in youth and adults. PLoS Biology, 20(12), e3001938.

Ke, J., Song, H., Bai, Z., Rosenberg, M. D., & Leong, Y. C. (2025). Dynamic brain connectivity predicts emotional arousal during naturalistic movie-watching. PLOS Computational Biology, 21(4), e1012994. 10.1371/journal.pcbi.1012994

Khazaei, S., Amin, Md. R., & Faghih, R. T. (2021). Decoding a Neurofeedback-Modulated Cognitive Arousal State to Investigate Performance Regulation by the Yerkes-Dodson Law. 2021 43rd Annual International Conference of the IEEE Engineering in Medicine & Biology Society (EMBC), 6551–6557. 10.1109/EMBC46164.2021.9629764

Knutson, B., Westdorp, A., Kaiser, E., & Hommer, D. (2000). FMRI Visualization of Brain Activity during a Monetary Incentive Delay Task. NeuroImage, 12(1), 20–27. 10.1006/nimg.2000.0593

Mahoney, J. J., Jackson, B. J., Kalechstein, A. D., De La Garza, R., & Newton, T. F. (2011). Acute, low-dose methamphetamine administration improves attention/information processing speed and working memory in methamphetamine-dependent individuals displaying poorer cognitive performance at baseline. Progress in Neuro-Psychopharmacology and Biological Psychiatry, 35(2), 459–465. 10.1016/j.pnpbp.2010.11.034

Martin, W. R., Sloan, J. W., Sapira, J. D., & Jasinski, D. R. (1971). Physiologic, subjective, and behavioral effects of amphetamine, methamphetamine, ephedrine, phenmetrazine, and methylphenidate in man. Clinical Pharmacology & Therapeutics, 12(2part1), 245–258. 10.1002/cpt1971122part1245

Miller, H. H., Shore, P. A., & Clarke, D. E. (1980). In vivo monoamine oxidase inhibition by d-amphetamine. Biochemical pharmacology, 29(10), 1347–1354.

Moeller, S. J., Honorio, J., Tomasi, D., Parvaz, M. A., Woicik, P. A., Volkow, N. D., & Goldstein, R. Z. (2014). Methylphenidate Enhances Executive Function and Optimizes Prefrontal Function in Both Health and Cocaine Addiction. Cerebral Cortex, 24(3), 643–653. 10.1093/cercor/bhs345

Molla, H., Keedy, S., DeBrosse, J., & De Wit, H. (2023). Methamphetamine enhances neural activation during anticipation of loss in the monetary incentive delay task. Cerebral Cortex Communications, 4(3), tgad014. 10.1093/texcom/tgad014

Molla, H., DeBrosse, J., Keedy, S., Lee, R., de Wit, H. (2025). Effects of methamphetamine on two measures of reward: Euphoria and neural activation to reward cues. Neuropsychopharmacology.

Nestor, L. J., Ghahremani, D. G., & London, E. D. (2023). Reduced neural functional connectivity during working memory performance in methamphetamine use disorder. Drug and Alcohol Dependence, 243, 109764. 10.1016/j.drugalcdep.2023.109764

O’Halloran, L., Cao, Z., Ruddy, K., Jollans, L., Albaugh, M. D., Aleni, A., Potter, A. S., Vahey, N., Banaschewski, T., Hohmann, S., Bokde, A. L. W., Bromberg, U., Büchel, C., Quinlan, E. B., Desrivières, S., Flor, H., Frouin, V., Gowland, P., Heinz, A., … Whelan, R. (2018). Neural circuitry underlying sustained attention in healthy adolescents and in ADHD symptomatology. NeuroImage, 169, 395–406. 10.1016/j.neuroimage.2017.12.030

Oswald, L. M., Wand, G. S., Wong, D. F., Brown, C. H., Kuwabara, H., & Brašić, J. R. (2015). Risky decision-making and ventral striatal dopamine responses to amphetamine: A positron emission tomography [11C]raclopride study in healthy adults. NeuroImage, 113, 26–36. 10.1016/j.neuroimage.2015.03.022

Riddle, E. L., Hanson, G. R., & Fleckenstein, A. E. (2007). Therapeutic doses of amphetamine and methylphenidate selectively redistribute the vesicular monoamine transporter-2. European Journal of Pharmacology, 571(1), 25–28. 10.1016/j.ejphar.2007.05.044

Rose, S. E., Janke, A. L., Strudwick, M. W., McMahon, K. L., Chalk, J. B., Snyder, P., & De Zubicaray, G. I. (2006). Assessment of dynamic susceptibility contrast cerebral blood flow response to amphetamine challenge: A human pharmacological magnetic resonance imaging study at 1.5 and 4 T. Magnetic Resonance in Medicine, 55(1), 9–15. 10.1002/mrm.20749

Rosenberg, M. D., Finn, E. S., Scheinost, D., Papademetris, X., Shen, X., Constable, R. T., & Chun, M. M. (2016a). A neuromarker of sustained attention from whole-brain functional connectivity. Nature Neuroscience, 19(1), 165–171. 10.1038/nn.4179

Rosenberg, M. D., Scheinost, D., Greene, A. S., Avery, E. W., Kwon, Y. H., Finn, E. S., Ramani, R., Qiu, M., Constable, R. T., & Chun, M. M. (2020). Functional connectivity predicts changes in attention observed across minutes, days, and months. Proceedings of the National Academy of Sciences, 117(7), 3797–3807. 10.1073/pnas.1912226117

Rosenberg, M. D., Zhang, S., Hsu, W.-T., Scheinost, D., Finn, E. S., Shen, X., Constable, R. T., Li, C.-S. R., & Chun, M. M. (2016b). Methylphenidate Modulates Functional Network Connectivity to Enhance Attention. The Journal of Neuroscience, 36(37), 9547–9557. 10.1523/JNEUROSCI.1746-16.2016

Shen, H., Wang, L., Liu, Y., & Hu, D. (2010). Discriminative analysis of resting-state functional connectivity patterns of schizophrenia using low dimensional embedding of fMRI. NeuroImage, 49(4), 3110–3121. 10.1016/j.neuroimage.2009.11.011

Shen, X., Tokoglu, F., Papademetris, X., & Constable, R. T. (2013). Groupwise whole-brain parcellation from resting-state fMRI data for network node identification. NeuroImage, 82, 403–415. 10.1016/j.neuroimage.2013.05.081

Silber, B. Y., Croft, R. J., Papafotiou, K., & Stough, C. (2006). The acute effects of d-amphetamine and methamphetamine on attention and psychomotor performance. Psychopharmacology, 187(2), 154–169. 10.1007/s00213-006-0410-7

Spencer, R. C., Devilbiss, D. M., & Berridge, C. W. (2015). The Cognition-Enhancing Effects of Psychostimulants Involve Direct Action in the Prefrontal Cortex. Biological Psychiatry, 77(11), 940–950. 10.1016/j.biopsych.2014.09.013

Stoy, M., Schlagenhauf, F., Schlochtermeier, L., Wrase, J., Knutson, B., Lehmkuhl, U., … & Ströhle, A. (2011). Reward processing in male adults with childhood ADHD—a comparison between drug-naive and methylphenidate-treated subjects. Psychopharmacology, 215, 467–481. 10.1007/s00213-011-2166-y

Swanson, J., Baler, R. D., & Volkow, N. D. (2011). Understanding the Effects of Stimulant Medications on Cognition in Individuals with Attention-Deficit Hyperactivity Disorder: A Decade of Progress. Neuropsychopharmacology, 36(1), 207–226. 10.1038/npp.2010.160

Van Den Heuvel, M. P., & Hulshoff Pol, H. E. (2010). Exploring the brain network: A review on resting-state fMRI functional connectivity. European Neuropsychopharmacology, 20(8), 519–534. 10.1016/j.euroneuro.2010.03.008

Volkow, N. D., Wang, G.-J., Fowler, J. S., Logan, J., Gerasimov, M., Maynard, L., Ding, Y.-S., Gatley, S. J., Gifford, A., & Franceschi, D. (2001). Therapeutic Doses of Oral Methylphenidate Significantly Increase Extracellular Dopamine in the Human Brain. The Journal of Neuroscience, 21(2), RC121–RC121. 10.1523/JNEUROSCI.21-02-j0001.2001

Vollenweider, F. X., Maguire, R. P., Leenders, K. L., Mathys, K., & Angst, J. (1998). Effects of high amphetamine dose on mood and cerebral glucose metabolism in normal volunteers using positron emission tomography (PET). Psychiatry Research: Neuroimaging, 83(3): 149–62. 10.1016/S0925-4927(98)00033-X

Wei, S., Xue, Z., Sun, W., Han, J., Wu, H., & Liu, X. (2021). Altered neural processing of reward and punishment in women with methamphetamine use disorder. Frontiers in Psychiatry, 12, 692266. 10.3389/fpsyt.2021.692266

